# Human eosinophil cationic protein manifests alarmin activity through its basicity and ribonuclease activity

**DOI:** 10.1101/2021.03.24.436724

**Authors:** Ayush Attery, Irene Saha, Prafullakumar Tailor, Janendra K. Batra

**Affiliations:** National Institute of Immunology, Aruna Asaf Ali Marg, New Delhi 110067, India; Department of Biochemistry, School of Chemical and Life Sciences, Jamia Hamdard, New Delhi 110062, India

**Author notes:** Department of Cell and Molecular Biology, Weizmann Institute of Science, Rehovot, Israel (Ayush Attery).

**Keywords:** Chemotaxis, EDN, ECP, Ribonucleases, alarmin

## Abstract

Eosinophil cationic protein (ECP), eosinophil derived neurotoxin (EDN), and human pancreatic ribonuclease (HPR) are members of the RNase A superfamily having similar catalytic residues and diverse functions. Alarmins are the endogenous mediators of innate immunity which activate or alarm the adaptive immune system by activating antigen presenting cells (APCs). EDN acts as an alarmin molecule and plays an important role in innate as well as adaptive immunity. EDN displays chemotactic activity for dendritic cells (DCs) and activates them, has antiviral and antiparasitic activities, and is rapidly released from immune cells. HPR only displays chemotactic activity while no such activity has been reported for ECP. In this study we show that ECP displays the chemotactic activity comparable to that of HPR and EDN. ECP also interacts with TLR-2 to activate NF-κB/AP-1 expression like EDN. The RNase activity of ECP, EDN and HPR, and basicity of ECP were found to be crucial determinants for their chemotactic activity for APCs, however for the DC maturation activity, RNase activity was not found to be essential. Bovine RNase A did not show any chemotactic activity despite having a very high RNase activity indicating that other determinants in addition to the RNase activity are involved in the chemotactic activity of ECP, EDN and HPR. The current study establishes that ECP also can act like an alarmin.

---

Eosinophils play an important role in clearing pathogens like bacteria, viruses and parasites, and contribute to innate immune defense (1–4). The anti-pathogenic activity of eosinophils is manifested through the eosinophil proteins, released during infection. These proteins include major basic protein (MBP), MBP-1 and MBP-2 present in the crystalloid core of the eosinophil granules, and eosinophil derived neurotoxin (EDN), eosinophil cationic protein (ECP) and eosinophil peroxidase (EPO) present in the matrix (5, 6). Of these granule proteins, EDN and ECP possess ribonuclease activity and have been classified as the members of RNase A superfamily (7). EDN has 67% sequence identity with ECP (8). However, despite being so similar ECP and EDN have differences in their catalytic and biological activities. Both ECP and EDN have been shown to have neurotoxic (9), antiviral (10) and antiparasitic activities (11–15). ECP in addition has cytotoxic (16, 17) and antibacterial activities (18).

Alarmins are endogenous mediators of immune system that activate immunity and protect hosts from exogenous danger signals (19, 20). To be called an alarmin, a molecule must be rapidly released from the cells in response to infection or tissue injury, it should act as a chemoattractant to antigen presenting cells (APCs) and activate them as well, and it should exhibit immune modulatory activities (20). EDN possesses all these activities and hence can be categorized as an alarmin. EDN has been shown to be rapidly released from multiple cell types like eosinophils, neutrophils and activated macrophages (21–24). EDN has been shown to selectively recruit dendritic cells (DCs) at the site of inflammation (25), activate them and produce antigen dependent Th2 immune response (26).

Chemotaxis, the phenomenon of migration towards a chemoattractant, plays an important role in immunity and maintaining homeostasis (27). It has been shown that EDN acts as a chemoattractant to DCs of both mouse and human, and its chemotactic activity towards DCs decreases in the presence of ribonuclease inhibitor (RI) in a dose-dependent manner suggesting that the RNase activity is crucial for the chemotactic activity (25). EDN stimulates DCs to release a variety of proinflammatory cytokines, chemokines, growth factors and soluble receptors (26). EDN and interestingly, human pancreatic ribonuclease (HPR) induced DC maturation by upregulating cell surface markers of DCs namely CD83, CD86 and MHC-II, switching cell surface markers from CCR5^+^ to CCR7^+^ and enhancing the capacity to stimulate allogenic T cell proliferation *in vitro* resulting in the release of proinflammatory cytokines and chemokines (22).

TLRs can induce an antimicrobial immune response by recognizing “non-self” components of foreign invaders (27, 28). Interaction of HPR and EDN with TLR2 of DCs leads to the activation of adaptive immunity. EDN has been shown to act through TLR2-MyD88 signalling pathway and activate NF-κB and MAPK in DCs (26). It has been demonstrated that EDN acts as an endogenous ligand specific to TLR2 independent of other TLRs (26).

ECP and EDN are very similar proteins; however it has not been investigated if ECP also has properties to be categorized as an alarmin like its counterpart, EDN. The antibacterial, cytotoxic and antiparasitic activities of ECP have been attributed to its highly basic nature and the unique basic residues, Arg22, Arg34, Arg61 and His64 (29).

EDN contains a stretch of nine amino acids, termed as loop L7, which is not present in HPR (Figure 1). ECP also contains loop L7 region, however, the composition of loop L7 in ECP is different than that in EDN. Three amino acids, Arg117, Pro120 and Gln122 in loop L7 of EDN differ between ECP and EDN (Figure 1). Mutations of these residues resulted in significant reduction in the antiviral activity of EDN, indicating them to be important for the interaction of EDN with the Respiratory Syncytial Virus (RSV) (30).

**Figure 1.**
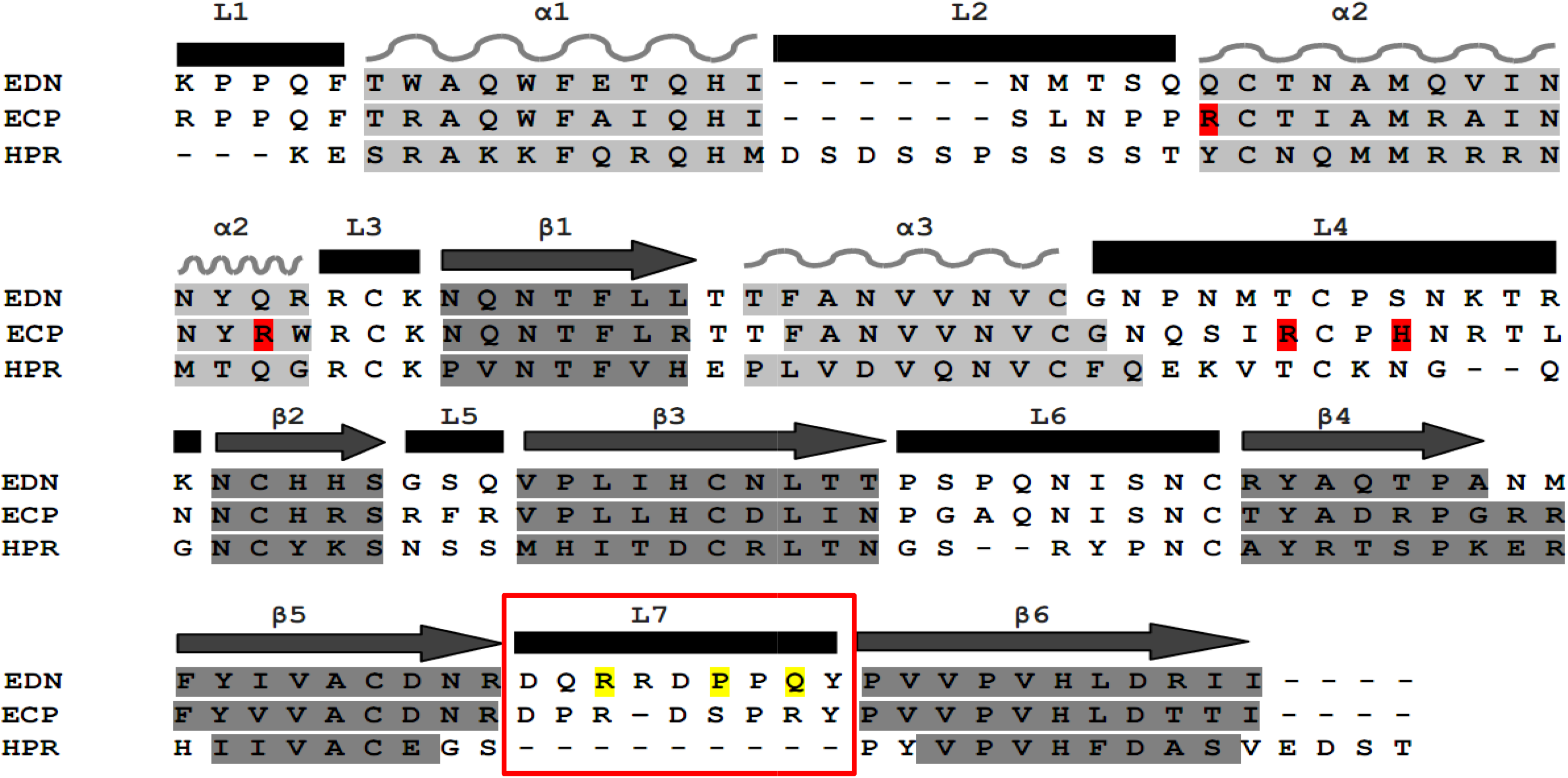
Multiple sequence alignment of human ribonucleases. The sequence of ribonucleases were taken from NCBI database and aligned by using multiple sequence alignment tool, Clustal-Omega. Secondary structural elements are shown in light grey (α-helix), dark grey (β-sheets) and black bars (loop regions). Residues in loop L7 of EDN and basic residues of ECP investigated in this study are highlighted in yellow and red respectively.

From the RNase A superfamily only EDN has been shown to manifest alarmin activity. In this study, we have carried out a comparative study of the chemotactic and DC maturation activities of ECP, EDN and HPR. Further, we have analyzed the roles of unique loop L7 in EDN, and basicity in ECP in their alarmin activity. We have also investigated the involvement of ribonuclease activity of these proteins in their chemotactic activity. The study demonstrates that ECP also manifests alarmin like activity, and the RNase activity of ECP, EDN and HPR is required for their DC maturation activity.

## EXPERIMENTAL PROCEDURES

### Protein expression and purification

Ribonucleases and their variants used in the current study are listed in Table 1. EDN has a unique secondary loop structure which is different from ECP and HPR. It is termed as loop L7 and three residues namely Arg117, Pro120 and Gln122 in this loop have been shown to be crucial for EDN’s biological activity (30). We have used three double mutants, EDN-R117A/P120A, EDN-P120A/Q122A, EDN-R117A/Q122A and a triple mutant, EDN-R117A/P120A/Q122A of these residues in L7 loop for our study. Also, as a control, we took EDN-H129A mutant of EDN which lacks RNase activity. In ECP, basic residues which are important for the antibacterial, antiviral and antiparasitic activities of ECP were identified earlier (29). These basic residues were mutated to alanine in combination and three variants of ECP, ECP-R22A/R34A, ECP-H64A/R34A, and ECP-H64A/R61A have been used in the current study (29). For HPR, we used an RNase inactive mutant, HPR-K41A as a control.

**Table 1.** Ribonuclease variants used in this study. The variants used in the current study were generated in our earlier studies and characterized for their other biological activities (29, 30).

| **Protein** | **Mutants** | **Importance** |
| --- | --- | --- |
| EDN | EDN-R117A/P120A<br>EDN-P120A/Q122A<br>EDN-R117A/Q122A<br>EDN-R117A/P120A/Q122A | Loop 7 mutants that show reduced anti-viral activity against RSV (Ref. 30). |
|  | EDN-H129A | RNase activity deficient mutant of EDN. |
| ECP | ECP-R22A/R34A<br>ECP-H64A/R34A<br>ECP-H64A/R61A | ECP mutants with reduced basicity that show reduced cytotoxic and antimicrobial activities of ECP (Ref: 29). |
|  | ECP-H15D | RNase activity deficient mutant of ECP. |
| HPR | HPR-K41A | RNase activity deficient mutant of HPR. |

All the recombinant proteins used in this study were expressed and purified from *E. coli* strain BL21-(λDE3)-RIL using clones that were in pVEX11 expression vector (29, 30). Bacterial cells transformed with appropriate plasmid were cultured in terrific broth supplemented with 100 μg/ml ampicillin. The cultures were induced with 0.5mM isopropyl-β-D-thiogalactopyaronide (IPTG) at A_600_ of 1.8 for 2 hours for HPR or 6 hours for ECP, EDN and their mutants. Cells were harvested by centrifugation at 4000xg for 15 minutes. All the proteins were found to accumulate in the cell in the form of inclusion bodies. The inclusion bodies were isolated, denatured and the recombinant proteins renatured as described by Buchner et al (31). Briefly, the inclusion bodies were dissolved in 6M guanidine hydrochloride, reduced with dithioerythritol, and renatured by diluting the protein 100-fold in a refolding buffer containing L-arginine-HCl and oxidized glutathione, and incubating at 10°C for 48 hrs for HPR (32) and 60 hrs for ECP and EDN (30). For HPR and EDN, dialysis was done for 12 hours in 20mM Tris-HCl buffer, pH 7.5 containing 100mM urea while for ECP it was in 20 mM sodium acetate buffer, pH 5.0 containing 100mM urea. The refolded proteins were then purified to homogeneity by successive cation exchange and gel filtration chromatography using SP Sepharose and Superdex 75 columns respectively. The purity of proteins was analyzed by SDS-PAGE. Proteins were characterized for their respective catalytic activity by RNase activity assay.

### Ribonuclease activity assay

The RNase activity of ECP, EDN, HPR and the mutants was assayed on yeast tRNA substrate following the method of Bond, 1988 (33). Briefly, an appropriate amount of each protein was incubated with 40μg of substrate in 10mM Tris-HCl, pH 7.5, at 37°C for an hour. The reaction was stopped by the addition of 5% (v/v) perchloric acid and 0.25% (w/v) uranyl acetate. The undigested large molecular weight RNA was precipitated on ice for 30 min and removed by centrifugation at 15000xg for 10 min. The acid soluble product, present in the supernatant, was quantified by measuring the absorbance at 260 nm.

### Dendritic cell isolation, purification and maturation

The DCs used in this study were isolated from mouse bone marrow (BMDCs) and cultured as described previously (34, 35). Briefly, bone marrow cells from femur were obtained by flushing femur with PBS supplemented with 2% heat inactivated FBS. These cells were then centrifuged and resuspended in Tris-ammonium chloride to lyse RBCs. The cells were spun and then filtered with a 70μm filter and were resuspended in RPMI-1640 supplemented with antibiotic and heat inactivated 10% FBS containing recombinant mouse GM-CSF (20ng/ml) and IL-4 (5ng/ml). Loosely adherent dendritic cells were collected and re-plated on day 7 in fresh RPMI medium and were harvested on day 8 and subjected to chemotaxis assay.

To analyse maturation, 0.5×10^6^ cells/well in a 24 well-plate were left untreated or treated with 10μg/ml each of ECP, EDN and HPR in separate wells for 48 hours at 37°C. After 48 hours, cells were stained with anti-mouse CD11c (BD Biosciences), CD80 (BD Biosciences), CD86 (eBioscience), CD40 (eBioscience) and MHCII (eBioscience), and analysed on FACS Canto II.

### Chemotaxis assay

Transwell inserts (Cornstar) of 5μm pore size and 6.5mm diameter size were used to check the chemotactic activity of ribonucleases. Briefly, 5μg/ml and 10μg/ml of proteins were incubated in chemotaxis medium, containing RPMI-1640 with 1% BSA, in the lower chamber and 100μl of chemotaxis medium were added in the insert well or upper chamber containing 1×10^6^ dendritic cells/well. The DCs were then allowed to migrate from upper chamber to lower chamber in a CO_2_ incubator for 90 minutes at 37°C. Cells were visualized microscopically under bright field microscope to check migration. After visual inspection, the transwells were removed and cells in the lower chamber were mixed and counted in a Neubauer’s haemocytometer. Wells with PBS alone were taken as blank.

### NF-κB and AP-1 expression

HEK-Blue™ hTLR2 cells are designed to study the expression of NF-κB by stimulation of TLR2. The expression of NF-κB and AP-1 was monitored using HEK Blue™-hTLR2 cells which express these proteins when TLR2 gets activated. These cells were obtained by co-transfection of the human TLR2 and SEAP (Secreted Embryonic Alkaline Phosphatase) genes into HEK293 cells. The SEAP reporter gene is placed under the control of the IFN-β minimal promoter fused to five NF-κB and AP-1-binding sites. Additionally, the CD14 co-receptor gene was transfected into these cells to enhance the TLR2 response. Stimulation with a TLR2 ligand activates NF-κB and AP-1 which induce the production of SEAP. Levels of SEAP can be easily determined with HEK-Blue™ Detection, a cell culture medium that allows for real-time detection of SEAP. The hydrolysis of substrate by SEAP produces a blue/purple color that can be visualized by naked eyes and can also be quantified spectrometrically by measuring absorbance at 655nm. Briefly, 5×10^4^ cells were incubated in HEK Blue detection media in the presence of 10, 25 and 50μg/ml of ribonuclease proteins. After 24 hours, these cells were visually monitored and absorbance at 655nm was measured spectrophometrically. The value of blank was subtracted from all the samples. Pam3 CSK4 was used as a positive control.

## RESULTS

### Multiple sequence alignment

The sequences of human ribonucleases, ECP, EDN and HPR were taken from NCBI database and were aligned by using multiple sequence alignment tool, Clustal Omega on EMBL server (Figure 1). The secondary structural elements, α-helix (light grey), β-sheets (dark grey) and loop regions (black bars) in these proteins are highlighted. Loop L7 of EDN and basic residues of ECP investigated in this study are highlighted in yellow and red respectively. Loop L7 region is shown in a box which is present in both ECP and EDN but absent in HPR.

### Expression, purification and ribonuclease activity of various proteins

Various proteins and their mutants used in this study are shown in Table 1. All proteins were expressed in *E. coli* and were localized in inclusion bodies from where they were isolated by denaturation and *in vitro* refolding. Proteins were purified by a two step purification strategy involving ion exchange chromatography and size exclusion chromatography. All the proteins were >90% pure, and single bands were observed on SDS-PAGE which corresponded to their respective molecular weights (data not shown).

The ribonuclease activity of human ribonucleases and their mutants was assessed using yeast tRNA as the substrate. HPR and EDN displayed higher RNase activity than ECP. EDN L7 loop double mutants, EDN-R117A/P120A, EDN-P120A/Q122A displayed similar activities as that of EDN, whereas EDN-R117A/Q122A and EDN-R117A/P120A/Q122A had negligible RNase activity (Table 2). All three basic residue mutants of ECP, ECP-R22A/R34A, ECP-H64A/R34A, and ECP-H64A/R61A were ribonucleolytically as active as the wild type protein. ECP-H15D, EDN-H129A and HPR-K41A are RNase inactive mutants of ECP, EDN and HPR respectively, which did not show any significant RNase activity (Table 2).

**Table 2.** Ribonuclease activity of ECP, EDN, HPR and their variants. Yeast tRNA was incubated with different concentrations of proteins. The undigested large molecular weight RNA was precipitated with perchloric acid and uranyl acetate on ice and removed by centrifugation. The acid soluble products were quantified by measuring the absorbance at 260 nm. Data represent the mean ± SEM of three independent experiments

| **Proteins** | **Specific Activity**<br>**(ΔAbsorbance /minute/mg protein)** | **% RNase activity** |
| --- | --- | --- |
| **ECP** | 780 ± 81 | 100 |
| ECP-R22A/R34A | 868 ± 34 | 111 |
| ECP-H64A/R34A | 682 ± 47 | 87 |
| ECP-H64A/R61A | 1107 ± 75 | 141 |
| ECP-H15D | 11 ± 6 | 1.4 |
| **EDN** | 11833 ± 739 | 100 |
| EDN-R117A/P120A | 10166 ± 514 | 86 |
| EDN-P120A/ Q122A | 13888 ± 732 | 117 |
| EDN-R117A/ Q122A | 482 ± 51 | 4 |
| EDN- R117A/P120A/Q122A | 466 ± 58 | 4 |
| EDN-H129A | 45 ± 16 | 0.4 |
| **HPR** | 20527 ± 650 | 100 |
| HPR-K41A | 37 ± 13 | 0.2 |

### ECP displays its chemotactic activity dependent on its catalytic activity and basicity

The chemotactic activity of ribonucleases and their mutants was assayed towards mouse DCs. ECP displayed chemotactic activity for DCs which was similar to that of EDN and HPR (Figure 2A). RNase A, used as a control showed a much lower activity (Figure 2A). Ribonucleolytically inactive mutants of ECP, EDN and HPR; ECP-H15D, EDN-H129A, and HPR-K41A displayed a significant reduction in their chemotactic activity indicating the RNase activity to be involved in the chemotactic activity of these proteins (Figure 2A).

**Figure 2.**
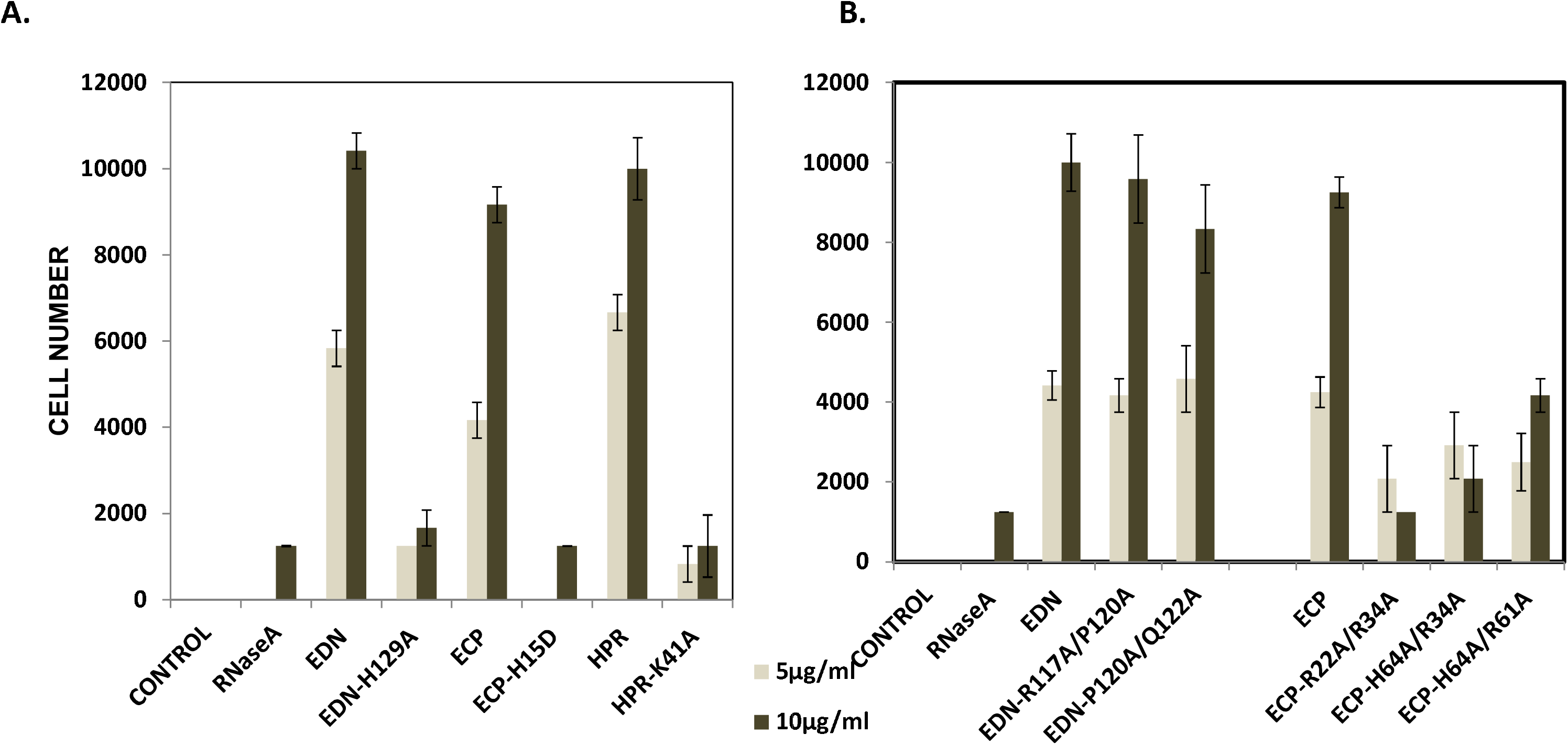
Chemotactic activity of human RNases and their mutants towards dendritic cells. Chemotactic activity towards DCs of RNase inactive mutants (A), and Loop L7 mutants of EDN and basic residue mutants of ECP (B) was assayed as described in the experimental section. Data represent mean ± SEM of two independent experiments done in triplicate.

The mutants of ECP with reduced basicity, ECP-R22A/R34A, ECP-H64A/R34A and ECP-H64A/R61A displayed reduced chemotactic activity towards DCs suggesting basicity to be an important determinant for ECP’s chemotactic activity (Figure 2B). RNase activity containing loop L7 mutants of EDN, EDN-R117A/P120A and EDN-P120A/Q122 also showed similar activity to that of EDN indicating loop L7 not to be critical for the chemotactic activity of EDN (Figure 2B).

### ECP induces dendritic cell maturation

Mouse GMFL-DCs were treated with various proteins and expression of CD40, CD80, CD86 and MHC-II was taken as the indicator of DC maturation. As compared to the control, there was an increase in the expression of CD40, and CD80 surface markers of CD11c^+^ DCs when they were treated with ECP, HPR and EDN indicating the maturation of DCs as a result of treatment with human RNases (Table 3). CD86 and MHC-II expression was also increased to moderate levels (Table 3). RNase A did not affect any of the markers indicating its inability to induce DC maturation.

**Table 3.** Effect of RNases on dendritic cell maturation. Effect of ECP, EDN, HPR and RNase A on CD11c^+^ dendritic cell maturation was analyzed by measuring expression of CD40, CD80, CD86 and MHC-II by flow cytometry. Briefly, 0.5×10^6^ cells were treated with 10 μg/ml of each of the proteins in separate wells for 48 hours at 37°C. At the end of incubation, cells were stained with anti-mouse CD11c, CD80, CD86, CD40 and MHCII antibodies and analysed on FACS Canto II. Values are given as percentage expression of markers on DCs and calculated by subtracting blank (untreated cells) from the test samples. The value for blank was 7.9, 19.7, 21.2 and 20.3 for CD40, CD80, MHCII and CD86 respectively.

| Proteins | % Expression |  |  |  |
| --- | --- | --- | --- | --- |
|  | CD40 | CD80 | MHC II | CD86 |
| ECP | 76.9 | 20.3 | 22.5 | 5.6 |
| EDN | 73.3 | 26.3 | 25 | 8 |
| HPR | 72.2 | 20.2 | 21.6 | 8.6 |

The loop L7 mutants of EDN, basic residue mutants of ECP and RNase activity deficient mutants of EDN, ECP and HPR also showed DC maturation activity, as indicated by higher surface expression of CD40 and CD80, like their respective wild type proteins (Table 4).

**Table 4.** Effect of RNase variants on dendritic cell maturation. Effect of variants of ECP, EDN and HPR on CD11c^+^ dendritic cell maturation was analyzed by measuring CD40 and CD80 expression as described in Table 3.

| Proteins | % Expression |  |
| --- | --- | --- |
|  | CD40 | CD80 |
| ECP | 81.6 | 42.5 |
| ECP-H15D | 80.5 | 59.1 |
| ECP-R22A/R34A | 64.2 | 67.7 |
| ECP-H64A/R61A | 73.9 | 66.1 |
| EDN | 86.3 | 53.5 |
| EDN-H129A | 83.5 | 49.1 |
| EDN-RPQ | 74.2 | 62.1 |
| HPR | 86.7 | 57 |
| HPR-K41A | 81.8 | 61.2 |

### ECP activates NF-κB and AP-1 expression via TLR2 signaling

HEK-Blue hTLR2 cells were used to investigate the effect of RNases on the level of NF-κB activation via TLR2. ECP, EDN and HPR and their mutants used in this study activated NF-κB expression in a dose-dependent manner in HEK-Blue hTLR2 cells indicating that they interacted with TLR2 (Figure 3A). Pam3-CSK4 was used as a positive control which displayed a high activity (Figure 3B). We have used recombinant LPS and recombinant flagellin as negative controls because they are known to interact with TLR4 and TLR5 respectively and hence must not display NF-κB expression. r-LPS and r-Flagellin indeed did not produce activation of NF-κB expression even at 10-fold higher concentration than Pam3 CSK4 (Figure 3B).

**Figure 3.**
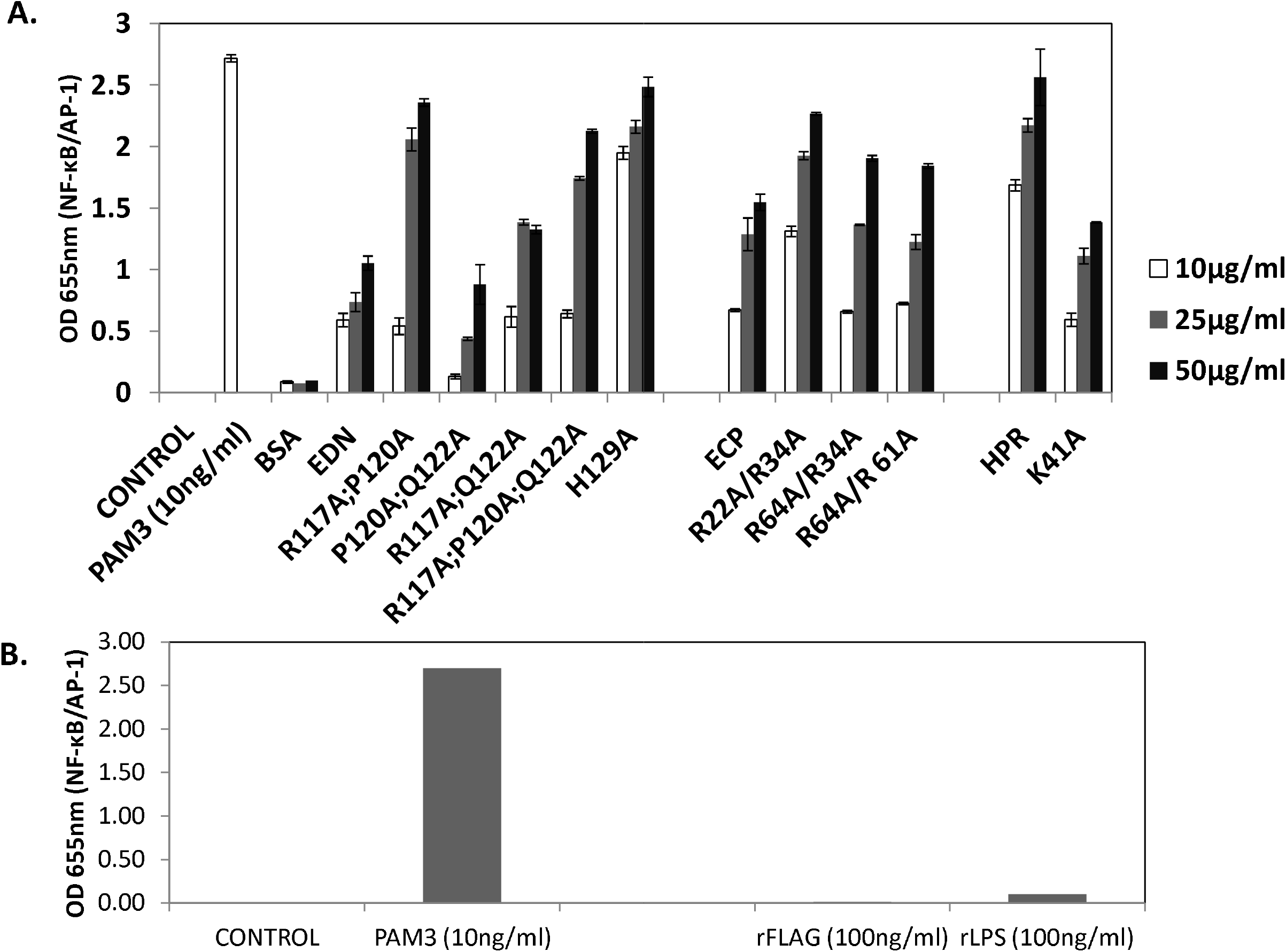
Effect of human RNases and their mutants on NF-κ B/AP-1 expression via TLR-2. Expression of NF-κB/AP-1 via TLR-2 was measured as described in the experimental section. **A.** Cells were treated with different concentrations of ribonucleases for 24 hours, **B.** TLR-2, TLR-5, and TLR-4 were stimulated by Pam3CSK-4, rFlagellin and rLPS respectively. Data represent the mean ± SEM of three independent experiments done in triplicate.

The NF-κB activation as a result of EDN, ECP, HPR and Pam3-CSK4 treatment was not affected by 4U/ml of ribonuclease inhibitor (Figure 4A). Treatment of ribonucleases with 10μg/ml of polymyxin B, which neutralizes LPS, also did not affect NF-κB expression suggesting that the observed effects are not due to LPS contamination in protein preparations (Figure 4B). DnaJ, a protein from *E. coli* used as a positively charged control protein did not activate NF-κB expression suggesting that the basicity alone is not a determinant for the activation of NF-κB expression (data not shown).

**Figure 4.**
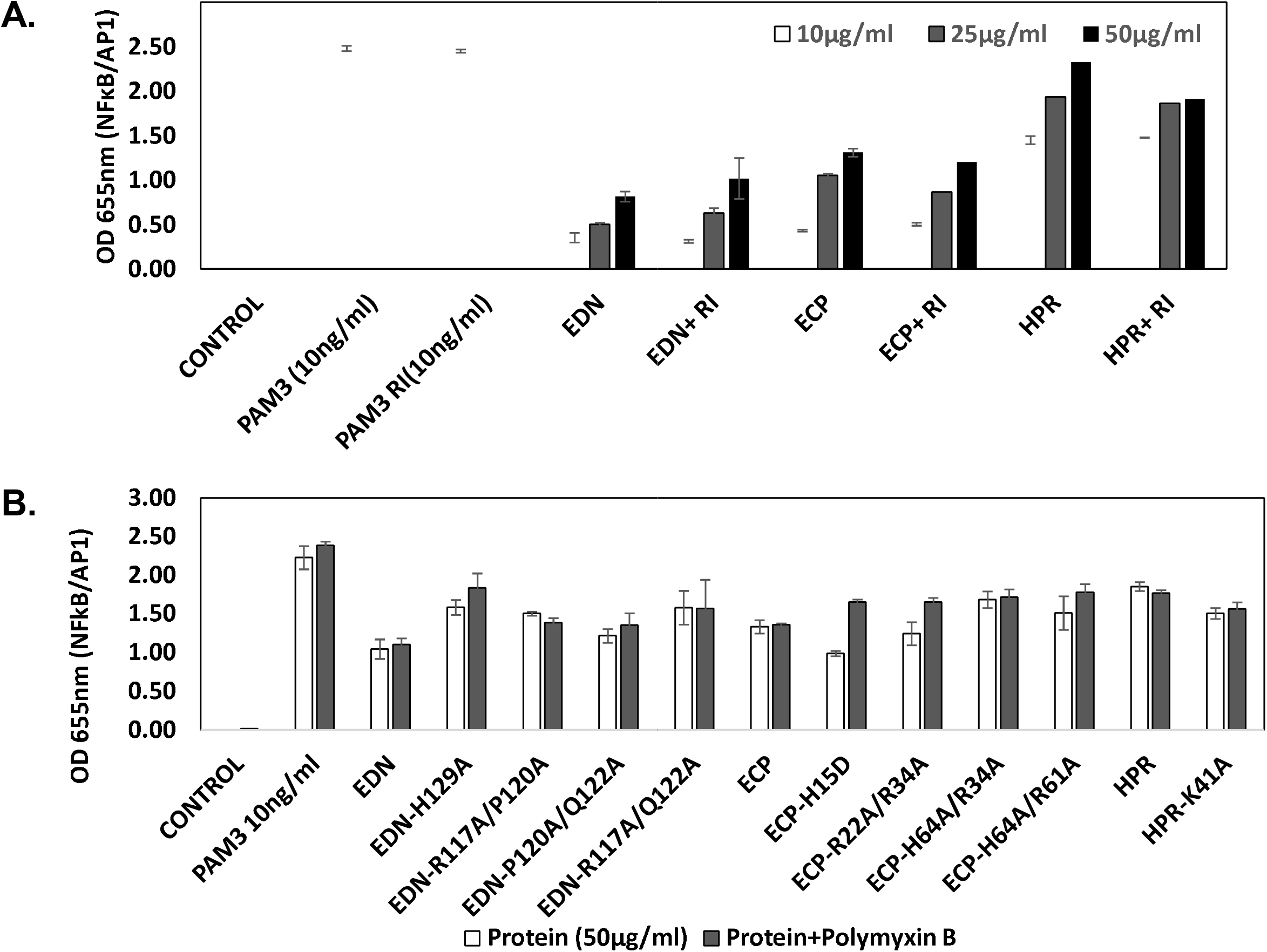
Effect of ribonuclease inhibitor and polymyxin B on NF-κ B/AP1 activation activity of RNases. Effect of RI on NF-κB/AP1 activation by RNase and mutants was assayed in the presence of ribonuclease inhibitor (A), and polymyxine B (B).

## DISCUSSION

Recruitment and activation of APCs towards the site of injury/stress is the most prominent feature of an alarmin molecule (20). EDN is known to act as an alarmin (25, 26). However, ECP has not been investigated for this activity. As mentioned previously, both EDN and HPR have the chemotactic activity for DCs (22). We show for the first time in this study that ECP also has this activity. It has been shown earlier that ribonuclease inhibitor inhibits EDN’s chemotactic activity (25). In this study, RNase inactive mutants of ECP, EDN and HPR did not show any significant activity demonstrating that RNase activity is indeed crucial for the chemotactic activity of these human RNases. Interestingly, RNase A did not show any such activity despite having a very high RNase activity, suggesting that there are determinants in addition to the RNase activity responsible for the chemotactic activity ofECP, EDN and HPR.

Loop L7 region of EDN is a unique determinant, and has been shown to be involved in the antiviral activity of EDN against RSV (30). We did not find any difference in the chemotactic and DC maturation activities of wild type and loop L7 mutants of EDN suggesting that loop L7 does not have any role in these activities of EDN.

Previously, we have shown that the positively charged amino acids unique to ECP, Arg22, Arg34, Arg61 and His64 are crucial for its cytotoxic, antibacterial and antiparasitic activities (29). We observed in this study that these basic amino acids also play a crucial role in the chemotactic activity of ECP. Thus, the current study demonstrates that the RNase activity of ECP, EDN and HPR, and basicity of ECP are crucial determinants for the chemotactic activity of these proteins towards APCs.

We demonstrate that all human ribonucleases used in this study stimulate maturation of DCs. Various known markers like CD40, CD80, CD86 and MHC-II were expressed on DCs after treatment with RNases. Loop L7 mutants of EDN and mutants of ECP with reduced basicity did not show any difference in DC maturation activity as compared to their wild type proteins suggesting that these determinants are not crucial for DC maturation activity of these proteins. For the DC maturation activity of these RNases, RNase activity was also not found to be essential. Further, the DC maturation activity was specifically limited to human RNases as bovine RNase A did not manifest this activity.

EDN activates MAPK signaling pathway (25) when it interacts with TLR-2 on dendritic cells which results in the activation NF-κB (26). We have observed that all the human ribonucleases, and their various mutants used in the current study interact with TLR-2 and activate NF-κB expression. DnaJ, a positively charged protein did not display the NF-κB activation suggesting that this activation is not a general activity of all basic proteins. Interaction of EDN, ECP and HPR with TLR-2 of DCs leads to the activation of adaptive immunity suggesting these ribonucleases to be at the intersection point of innate and adaptive immunity.

In conclusion, the current study demonstrates that like EDN, ECP also can act like an alarmin. ECP also interacts with TLR-2 to activate NF-κB/AP-1 expression. The RNase activity of HPR, ECP and EDN, and basicity of ECP were found to be crucial determinants for chemotactic activity of these proteins towards APCs, however for the DC maturation activity, RNase activity was not found to be essential. As bovine RNase A did not show any chemotactic activity despite having a very high RNase activity, it appears there are determinants in addition to the RNase activity for the chemotactic activity of ECP, EDN and HPR.

## ACKNOWLEDGEMENTS

This work was supported by grants to the National Institute of Immunology, New Delhi from the Department of Biotechnology, Government of India. AA thanks the Council of Scientific and Industrial Research, India for a senior research fellowship.

